# Multi-species mechanistic model of functional response provides new insights into predator-mediated interactions in a vertebrate community

**DOI:** 10.1101/2021.10.22.464289

**Authors:** Andréanne Beardsell, Dominique Gravel, Jeanne Clermont, Dominique Berteaux, Gilles Gauthier, Joël Bêty

## Abstract

Prey handling processes are considered a key driver of short-term positive indirect effects between prey sharing the same predator. However, a growing body of research indicates that predators are rarely limited by such processes in the wild. Density-dependent changes in predator foraging behavior can also generate positive indirect effects but they are rarely included as explicit functions of prey densities in functional response models. With the aim of untangling proximate drivers of species interactions in natural communities and improve our ability to quantify interaction strength, we extended the Holling multi-species model by including density-dependent changes in predator foraging behavior. Our model, based on species traits and behavior, was inspired by the vertebrate community of the arctic tundra, where the main predator (the arctic fox) is an active forager feeding primarily on cyclic small rodent (lemming) populations and eggs of various tundra-nesting bird species. Short-term positive indirect effects of lemmings on birds have been documented over the circumpolar Arctic but the underlying proximate mechanisms remain poorly known. We used a unique data set, containing high-frequency GPS tracking, accelerometer, behavioral, and experimental data to parameterize the multi-species model, and a 15-year time series of prey densities and bird nesting success to evaluate interaction strength between species. Our results showed that: (i) prey handling processes play a minor role in our system and (ii) density-dependent changes in predator foraging behavior can be the proximate drivers of a predominant predator-mediated interaction observed in the arctic tundra. Mechanisms outlined in our study have been little studied and may play a significant role in natural systems. We hope that our study will provide a useful starting point to build mechanistic models of predation, and we think that our approach could conceivably be applied to a broad range of food webs.

## 2. Introduction

Interaction between predator and prey is a complex phenomenon to represent in multi-species natural communities. Various mathematical models are used to quantify prey acquisition by the predator and to investigate the nature and the strength of species interactions within food webs (Abrams et al., 1998; Pawar et al., 2012; Baudrot et al., 2016; Chan et al., 2017). Several studies have compared how a predator acquisition rate varies with prey density using statistical approaches (reviewed by Novak and Stouffer 2020) but few of them tackle the underlying mechanisms explicitly. While statistical approaches can faithfully reproduce empirical observations, they do not give clear insights into the mechanisms underpinning interaction rates. Moreover, the sample size is often insufficient to obtain the statistical power needed to properly discriminate models (Novak and Stouffer, 2020). Process-based mechanistic models (hereafter referred to as a mechanistic model) may help in untangling proximate drivers of species interactions (Connolly et al., 2017; Griffen, 2021) and can improve our ability to adequately quantify the strength of interactions in natural communities (Spalinger and Hobbs, 1992; Beardsell et al., 2021; Delong et al., 2021).

Holling multi-species model is often used to model predation rates in multi-prey systems (e.g., Brose et al. 2005; McLellan et al. 2010; Barraquand et al. 2015; Serrouya et al. 2015). The summation of the handling time of all prey items is a critical component of this equation and can generate indirect interactions among prey. Increasing abundance of one species saturates the predator because of limited prey handling capacities (which includes chasing, killing and eating) and thereby indirectly releases predation pressure on other prey. This equation is often at the core of more complex food web models (Schneider et al., 2016; Tyson and Lutscher, 2016; Barrios-O’Neill et al., 2019) and handling time is considered a key potential driver of short-term positive effects observed among prey (Abrams and Matsuda, 1996; Abrams et al., 1998). However, the role of handling processes in predator-mediated interactions lacks definitive evidence in the wild and a growing body of research indicates that predators are rarely limited by handling processes at the highest prey densities observed in natural systems (Jeschke et al., 2002; Novak, 2010; Chan et al., 2017; Preston et al., 2018; Beardsell et al., 2021).

It is unlikely that predators simply acquire more prey as they increase in abundance. Non-linearities in functional responses are susceptible to influence predation rates via several mechanisms. For instance, the density of a prey may modulate the predator state (e.g., hunger level, reproductive status), which in turn could have an impact on other prey consumed by that predator. The challenge for ecologists is thus to evaluate and properly represent how the acquisition rate is influenced not only by the density of a given prey item as in typical Holling type II functional responses, but also by variation in abundance of the predator and all other potential prey items. Density-dependence has been considered in several functional response models (Hassell et al., 1977; Abrams, 1982) but rarely as explicit functions of prey densities (Stouffer and Novak, 2021). Such density-dependent changes in predator foraging behavior can generate positive effects between prey (Abrams and Matsuda, 1996, 1993) and warrant additional attention in natural predator-prey systems (Stouffer and Novak, 2021).

Our objectives are twofold. First, we extend the Holling multi-species model to develop a mechanistic model of acquisition rates that includes prey density-dependent changes in both predator handling time, and foraging behavior. Second, we illustrate this model using the predator-prey dynamics of a multi-species system in the arctic tundra to identify the proximate drivers of the well-known short-term positive indirect effects of cyclic rodents on nesting birds (see below). This type of predator-mediated effects is widespread between prey sharing a predator and can affect species abundance and coexistence in various ecosystems (Bonsall and Hassell, 1997; Duchesne et al., 2021). We thus evaluate three hypotheses that could explain such indirect interactions, motivated by theory and extensive empirical observations (Table 1 and Fig. 1). The first hypothesis is based on the Holling multi-species model, in which prey handling processes reduce the time available for searching other prey and result in positive indirect effects between prey. The second and the third hypothesis extend the Holling multi-species model to include prey density-dependence in prey handling time, and in predator foraging behavior, respectively (Table 1).

**Table 1.**
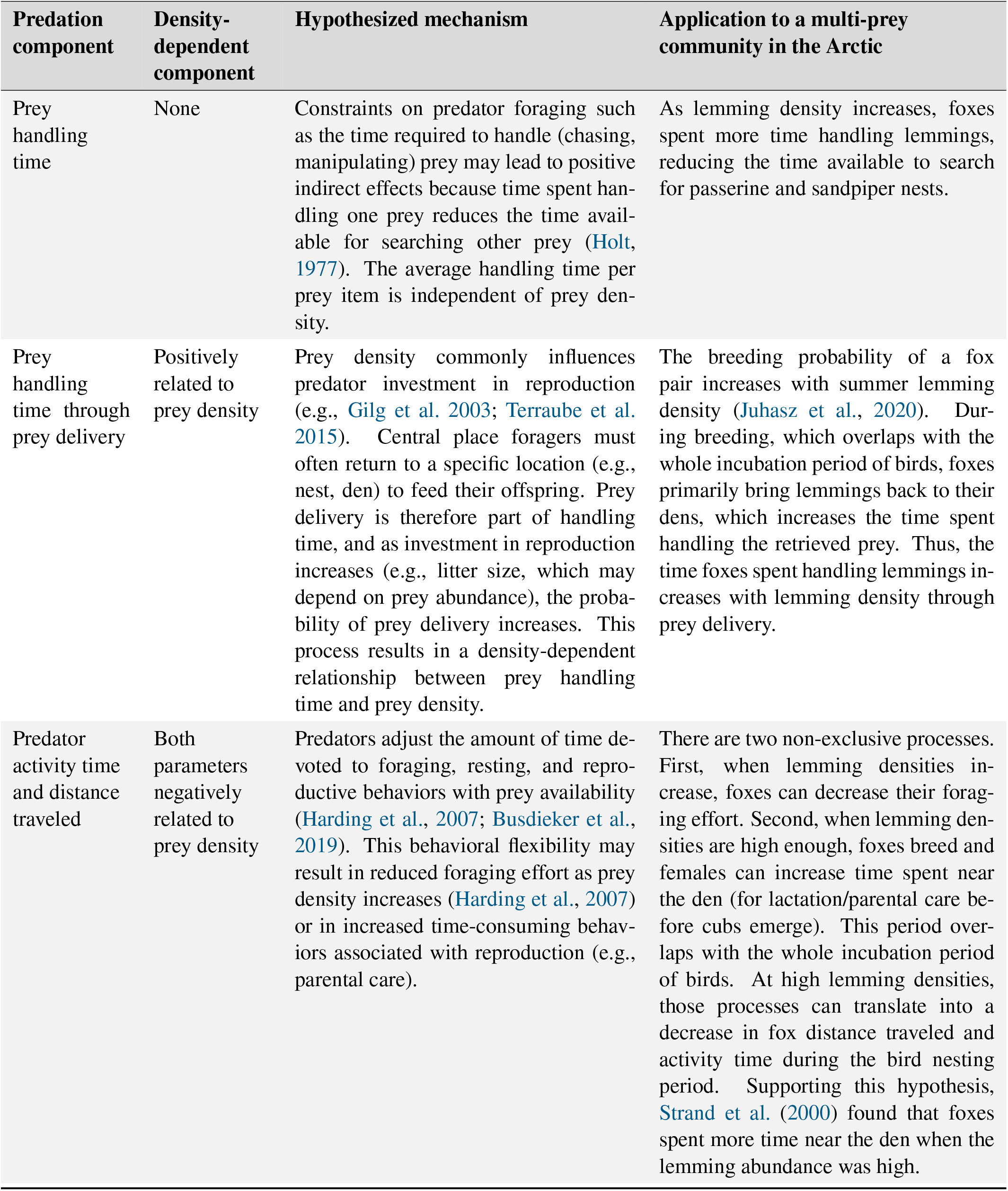
Three hypothesized mechanisms underlying the short-term positive effects of a cyclic prey (prey 1; lemmings) on two prey species (prey 2 and 3; passerine and sandpiper nests, respectively) through a common predator (arctic fox).

| Predation component | Density-dependent component | Hypothesized mechanism | Application to a multi-prey community in the Arctic |
| --- | --- | --- | --- |
| Prey handling time | None | Constraints on predator foraging such as the time required to handle (chasing, manipulating) prey may lead to positive indirect effects because time spent handling one prey reduces the time available for searching other prey (Holt, 1977). The average handling time per prey item is independent of prey density. | As lemming density increases, foxes spent more time handling lemmings, reducing the time available to search for passerine and sandpiper nests. |
| Prey handling time through prey delivery | Positively related to prey density | Prey density commonly influences predator investment in reproduction (e.g., Gilg et al. 2003; Terraube et al. 2015). Central place foragers must often return to a specific location (e.g., nest, den) to feed their offspring. Prey delivery is therefore part of handling time, and as investment in reproduction increases (e.g., litter size, which may depend on prey abundance), the probability of prey delivery increases. This process results in a density-dependent relationship between prey handling time and prey density. | The breeding probability of a fox pair increases with summer lemming density (Juhasz et al., 2020). During breeding, which overlaps with the whole incubation period of birds, foxes primarily bring lemmings back to their dens, which increases the time spent handling the retrieved prey. Thus, the time foxes spent handling lemmings increases with lemming density through prey delivery. |
| Predator activity time and distance traveled | Both parameters negatively related to prey density | Predators adjust the amount of time devoted to foraging, resting, and reproductive behaviors with prey availability (Harding et al., 2007; Busdieker et al., 2019). This behavioral flexibility may result in reduced foraging effort as prey density increases (Harding et al., 2007) or in increased time-consuming behaviors associated with reproduction (e.g., parental care). | There are two non-exclusive processes. First, when lemming densities increase, foxes can decrease their foraging effort. Second, when lemming densities are high enough, foxes breed and females can increase time spent near the den (for lactation/parental care before cubs emerge). This period overlaps with the whole incubation period of birds. At high lemming densities, those processes can translate into a decrease in fox distance traveled and activity time during the bird nesting period. Supporting this hypothesis, Strand et al. (2000) found that foxes spent more time near the den when the lemming abundance was high. |

**Fig. 1.**
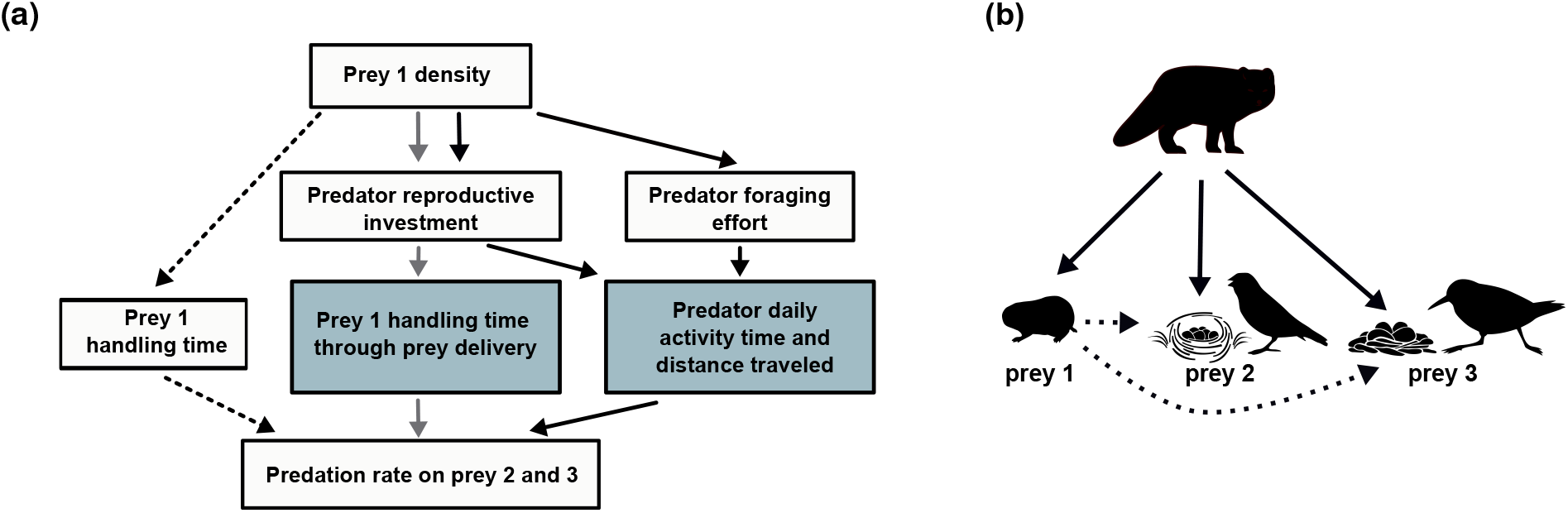
(a) Schematic representation of three hypothesized mechanisms underlying the short-term positive effect of a cyclic prey (prey 1, lemmings) on two prey species (prey 2 and 3, passerine and sandpiper nests, respectively) through a common predator (arctic fox). Different arrows (dotted, grey and black) correspond to each hypothesis. The blue boxes indicate the parameters where density-dependence was included. See Table 1 for a detailed description of each hypothesis. (b) Diagram of a simplified arctic food web indicating the direct (solid arrows) and indirect (dashed arrows) links between the predator (arctic fox), prey 1 (lemmings), prey 2 (passerine nests) and 3 (sandpiper nests).

We develop the multi-species mechanistic model for the arctic fox, an active-searching top predator of the arctic tundra that feeds primarily on lemmings, as well as on bird eggs during summer (Angerbjörn et al., 1999; Giroux et al., 2012). Within this predator-prey system, lemmings are the most abundant prey and show population cycles with a maximum density occurring every 3-5 years (Fauteux et al., 2015). Fox predation pressure on eggs of ground-nesting birds is generally released when lemming density is high, leading to short-term positive indirect effects of lemmings on bird nesting success (Summers et al., 1998; Nolet et al., 2013; McKinnon et al., 2014). This classic example of predator-mediated effects among vertebrates was studied across the circumpolar arctic but the underlying proximate mechanisms remain unknown (Underhill et al., 1993; Summers et al., 1998; Nolet et al., 2013; McKinnon et al., 2014). We parameterize the model using a combination of behavioral, demographic, and experimental data acquired over 20 years in the high-arctic tundra. As recent single-prey mechanistic model indicates that arctic fox is not limited by handling processes at the highest lemming densities observed in our study system (Beardsell et al., 2021), we expect density-dependent changes in the predator foraging behavior to be the main proximate driver of the short-term positive effects of lemmings on arctic bird nesting success.

## 3. Methods

### 3.1 Study system

The mechanistic model of multi-prey functional response was developed using data from a long-term ecological monitoring on Bylot Island, Nunavut, Canada (73° N; 80° W). Two cyclic species of small mammals are present, including the brown (*Lemmus trimucronatus*) and collared (*Dicrostonyx groenlandicus*) lemmings. Ground-nesting birds present include passerines (mostly Lapland longspur, *Calcarius lapponicus*) and sandpipers (primarily Baird’s Sandpiper (*Calidris bairdii*) and White-rumped Sandpiper (*Calidris fuscicollis*)). The monitoring area of lemmings, passerine and sandpiper nests is located within the Qarlikturvik Valley (72°85’N, 78°85’W). Sandpipers and passerines nest at relatively low densities (2 and 7 nests km^-2^ on average, respectively). During the breeding season (June until July) passerines and sandpipers lay an average of 5 and 4 eggs, which they incubate for 12 and 21 days, respectively (Gauthier et al., 2013; McKinnon et al., 2014; Hussell and Montgomérie, 2020). Sandpiper chicks typically leave the nest within 24 hours of hatching (McKinnon et al., 2014), while passerine chicks remain in the nest for 9-10 days (Hussell and Montgomérie, 2020).

The arctic fox is the main egg predator of ground-nesting birds on Bylot Island (McKinnon and Bêty, 2009; Royer-Boutin, 2015). The incubation period for birds (typically mid-June to early mid-July) overlaps with the lactation period for foxes. Fox gestation period is around 52 days and births usually occur in late May on Bylot Island (Audet et al., 2002; Morin, 2015). Fox cubs are weaned after 6-7 weeks (Audet et al., 2002). Arctic foxes maintain summer territories (averaging 10 km^2^) with little overlap (Grenier-Potvin et al., 2021), which limits interference between foxes within territories. Like many other animals (Vander Wall, 1990), arctic foxes generally predate more prey than they immediately consume (food hoarding behavior) and thus hide a large proportion of the prey they capture (Samelius and Alisauskas, 2000; Careau et al., 2007).

The monitoring area of passerines and sandpipers nest is located ~30 km away from a colony of greater snow geese (*Anser caerulescens atlanticus*), which is concentrated in an area of 50-70 km^2^ (McKinnon et al., 2014). We thus excluded geese from the multi-species model since they are virtually absent in the monitoring area. Isotopic studies confirmed that the contribution of goose eggs to the diet of foxes in the monitoring area was limited (Giroux et al., 2012). However, fox movement data (GPS and accelerometer; see details below) used to parameterize the model were collected within the boundary of the goose colony. Snow geese can influence fox habitat selection and diet within the colony (Giroux et al., 2012; Lamarre et al., 2017). Such effects could slightly bias the average values of a few parameters used in our models (e.g., daily distance traveled, time spent active). However, we are highly confident that the effect of lemming fluctuations on fox behavior is relatively similar across the landscape because fox reproduction and predation pressure on bird nests are strongly influenced by lemming density both inside and outside the goose colony (Giroux et al., 2012; Lamarre et al., 2017; Duchesne et al., 2021).

### 3.2 Deriving mechanistic models of multi-species functional response

#### 3.2.1 No density-dependence (model A)

We used the Holling multi-species model as a starting point to build the multi-species functional response model:

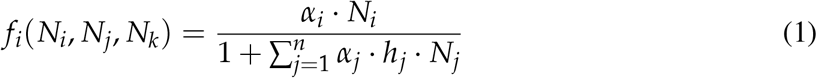

Here, *f_i_* is the acquisition rate of the predator on a prey species (*i*) as a function of all prey densities (*N_i_*, *N_j_* and *N_k_*). Parameter *α_i_* is the capture efficiency (km^2^ day^-1^) of the predator, whereas *h_i_* is the handling time for prey type *i* (day prey item^-1^). Capture efficiency is obtained by the product of the daily distance traveled by the predator (s; km day^-1^), reaction distance (*d_i_*; km), detection (*z_i_*) and attack probabilities (*k_i_*) of the prey by the predator, and the success probability (*p_i_*) of an attack (Table 2). *s* is defined as the daily distance traveled when the predator is active while *d_i_* is defined as the maximum distance at which the predator and prey *i* can detect or react to each other (in 2D, detection region = 2*d*; Pawar et al. 2012). Handling time (*h_i_*) includes the time spent chasing a prey *i* once attacked (*T_ci_*) and the time spent manipulating a prey *i* once captured (*T_mi_*):

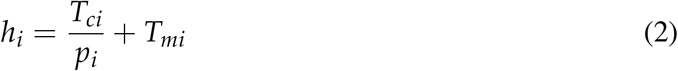

**Table 2.** Values of parameters used in the multi-species functional response model of a mammalian predator (arctic fox) to density of a cyclic prey (prey 1; lemmings) and two other prey species (prey 2 and 3; passerine and sandpiper nests, respectively). Parameter values were estimated from a combination of high-frequency GPS and accelerometry (23 summer foxes, 2018-2019), behavioral observations (*n* = 124 hours, 1996-2019) and camera traps (2006-2016) data.

| Parameter name | Symbol | Value(s) | Unit |
| --- | --- | --- | --- |
| Proportion of time spent active in a day | $\beta$ | 0.5 | - |
| Daily distance traveled (when $\beta = 0.5$) | $s$ | 41 | km day<sup>-1</sup> |
| **Lemmings** |  |  |  |
| Lemming density | $N_1$ | 0-700 | ind. km<sup>-2</sup> |
| Maximum reaction distance | $d_1$ | 0.0075 | km |
| Average detection and attack probability within the reaction distance | $z_1 * k_1$ | 0.15 | - |
| Success probability | $p_1$ | 0.51 | - |
| Chasing time | $T_{c1}$ | $1.0 \times 10^{-3}$ | day ind.<sup>-1</sup> |
| Consumption time | $T_{e1}$ | $3.8 \times 10^{-4}$ | day ind.<sup>-1</sup> |
| Consumption probability | $e_1$ | 0.48 | - |
| Hoarding time | $T_{o1}$ | $4.9 \times 10^{-4}$ | day ind.<sup>-1</sup> |
| Hoarding probability | $o_1$ | 0.32 | - |
| Delivering time | $T_{d1}$ | $3.9 \times 10^{-3}$ | day ind.<sup>-1</sup> |
| Delivering probability | $de_1$ | 0.20 | - |
| **Passerine nests** |  |  |  |
| Passerine nest density | $N_2$ | 0-15 | nests km<sup>-2</sup> |
| Maximum reaction distance | $d_2$ | 0.02 | km |
| Average detection probability within the reaction distance | $z_2$ | 0.12 | - |
| Consumption time | $T_{e2}$ | $3.6 \times 10^{-4}$ | day nest<sup>-1</sup> |
| **Sandpiper nests** |  |  |  |
| Sandpiper nest density | $N_3$ | 0-7 | nests km<sup>-2</sup> |
| Maximum reaction distance | $d_3$ | 0.085 | km |
| Average detection probability within the reaction distance | $z_3$ | 0.029 | - |
| Consumption time | $T_{e3}$ | $2.8 \times 10^{-3}$ | day nest<sup>-1</sup> |

In equation 1, time spent searching by the predator is only reduced by prey handling time. Although it has been recognized that predator activity time is an important component of the functional response (Abrams, 1982, 1990), this parameter is rarely included in functional response models. We therefore extend equation 1 to explicitly include the proportion of time that the predator spent active (Appendix S1; Eq. S4). The total time available in a day is now reduced by i) *β* which expresses the proportion of time spent active in a day by ii) 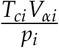 which is the time spent chasing a prey *i* once encountered and by iii) *T_mi_V_αi_* which is the time spent handling a prey *i* if subdued. The model including *β* takes the following form:

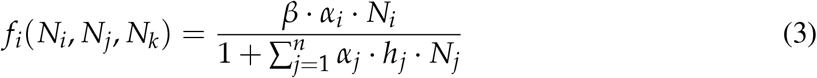

Equation 3 is the basic model, without density dependence in the predation components. An equivalent equation of the predator acquisition rate on prey *j* and *k* can be obtained by substituting all *i*’s for *j*’s (or *k*) and vice versa. The complete model derivation is available in Appendix S1. Below, we provide details on the inclusion of prey density-dependence in models B and C.

#### 3.2.2 Density-dependence in prey *i* handling time (model B)

During the breeding season, central-place foragers must often return to a specific breeding location (e.g., den, nest) to feed their offspring. This behavior is likely to increase the handling time for prey brought back to the breeding location. Prey delivery is therefore part of handling time, and as investment in reproduction increases, prey delivery increases until reaching a plateau. If prey *i* are returned to the den, prey *i* handling time can be expressed as a function of prey *i* density (N_i_) as follows:

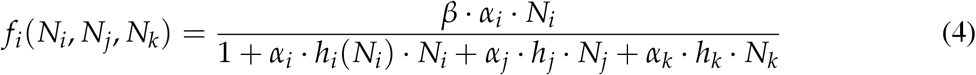

#### 3.2.3 Density-dependence in predator activity time and distance traveled (model C)

Predators can adjust the time devoted to foraging, resting, and reproductive behaviors with prey availability (Harding et al., 2007; Busdieker et al., 2019). This behavioral flexibility may result in reduced foraging effort as prey density increases (Harding et al., 2007) or in increased time-consuming behaviors associated with reproduction (e.g., parental care). Thus, this translates into explicit relationships between distance traveled (s), proportion of time spent active (*β*), and prey density (in the equations, prey *i*):

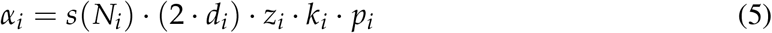

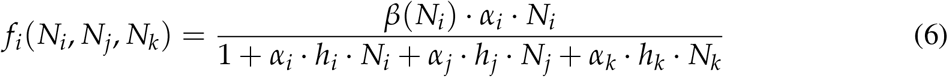

#### 3.2.4 Application of the functional response model to a multi-prey community in the Arctic

Equation 3 was adapted to each prey species according to their anti-predator behavior and the fox behavior observed during the bird nesting season (Beardsell et al., 2021). Figure 2 provides an overview of the mechanistic model adapted to each prey species (*i* = 1 for lemmings, *j* = 2 for passerine nests and *k* = 3 for sandpiper nests).

**Fig. 2.**
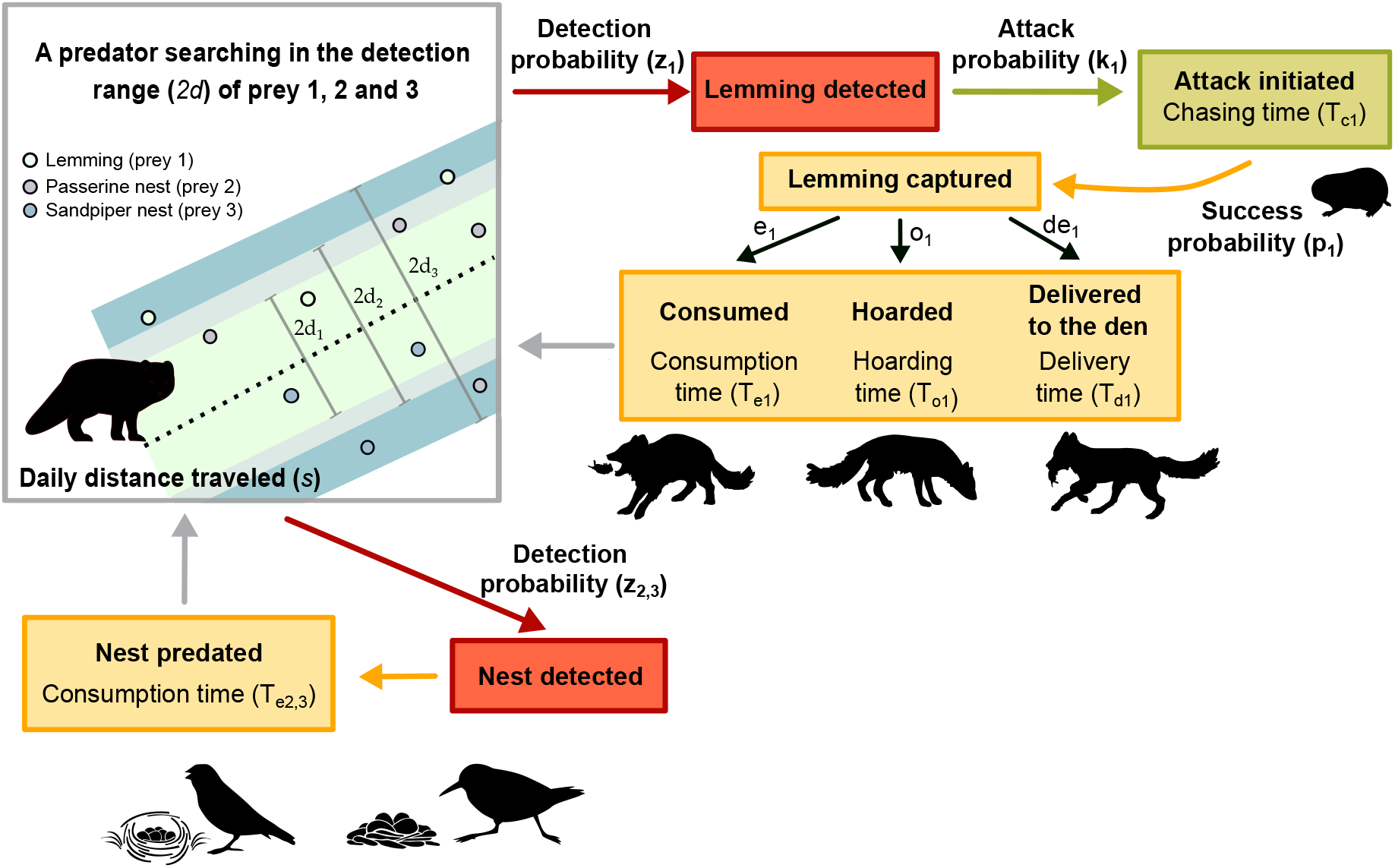
Conceptual mechanistic model of predator (arctic fox) functional response to density of a cyclic prey (prey 1; lemmings) and other prey species (prey 2 and 3; ground-nesting birds). Each box represents a component of predation (search, detection, attack and predation per se). Arrows represent the probability that the predator reaches the next component. When there is no parameter near the arrow, the probability to reach the next component is assumed to be 1. The model represented has no density dependence in the parameters.

As passerines and sandpipers are unable to protect their clutches against arctic foxes (Smith and Edwards, 2018; Hussell and Montgomérie, 2020), their nests are always predated when detected (no chasing time and the attack probability of the nest when detected is 1) and consumed immediately (Beardsell et al., 2021). Thus, the handling time of passerine and sandpiper nests (*h*_2_ and *h*_3_) includes only nest consumption time (*T_e2_* and *T_e3_*).

The manipulation time of lemmings is divided in three mutually exclusive behaviors: the lemming is either 1) consumed, 2) hidden, or 3) transported (Careau et al., 2007). The handling time of lemmings (*h*_1_) is hence expressed as follows:

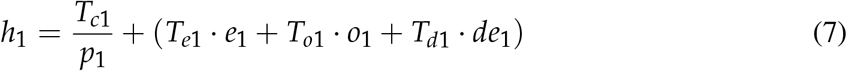

where *e*_1_, *o*_1_ and *d*_*e*1_ are respectively the probability that a lemming is consumed, hoarded or delivered to the den. The sum of *e*_1_, *o*_1_ and *d*_*e*1_ equals 1. *T*_*e*1_, *T*_*o*1_ and *T*_*d*1_ are the time associated with each behavior. As many other carnivores (Jeschke, 2007), arctic foxes probably often meet or exceed their daily energy requirements during the summer. However, there is no evidence that short-term energy needs influence arctic foxes behavior since they can harvest more prey than they consume in the short term and prey are cached for later consumption (e.g., during winter; Careau et al. 2007). Thus, prey digestion time was not included in the model.

### 3.3 Parameter values

From June to August 2018 and 2019, 16 foxes (7 females and 9 males) were fitted with high-frequency GPS collars and tri-axial accelerometers (95 g, 2.6–3.3% of body mass; Radio Tag-14, Milsar, Romania) to monitor their movements and behaviors. Of these, 7 were equipped in both years, for a total of 23 summer foxes (8 foxes in 2018 and 15 in 2019). Foxes were captured using cage traps (Tomahawk Live Trap Company, USA) or Softcatch #1 padded leghold traps (Oneida Victor Inc. Ltd., USA). GPS fix intervals were set to 4 minutes (360 fixes day ^-1^) and the location error was 11 m (Poulin et al., 2021); 30-second bursts of accelerometry data were collected every 4.5 minutes at 50 Hz (320 bursts day^-1^; Clermont et al. manuscript under review). We extracted foxes’ daily activity budget from accelerometry data (Clermont et al. manuscript under review). We estimated the proportion of time spent active by subtracting the proportion of time spent resting from 1. We estimated an average proportion of time spent active using a linear mixed model with year and individual-fox as random effects. The average proportion of time spent active in a day (*β*) was 0.50 (*n* = 371 fox-days; 95% CI (0.40-0.60); Table 2) and ranged from 0.29 to 0.64.

Distance traveled by foxes each day (km day^-1^) was estimated by adding linear distances between successive GPS locations and was extracted from 5 June to 9 July to cover the incubation period of most birds. Days with less than 75% of observations (i.e., 270 daily fixes) were excluded from analyses to avoid underestimating daily distances. Since the distance and the proportion of time spent active were closely correlated (Fig. 3a), we applied a linear mixed model to estimate the average daily distance with the distance as the response, the proportion of time spent active as a fixed factor, and individual-fox and year as random effects. The predicted average daily distance traveled was 41 km (*n* = 371 fox-days; 95% CI (32-49 km)) and ranged from 19 to 62 km while setting the proportion of time spent active at the average (i.e., 0.5).

**Fig. 3.**
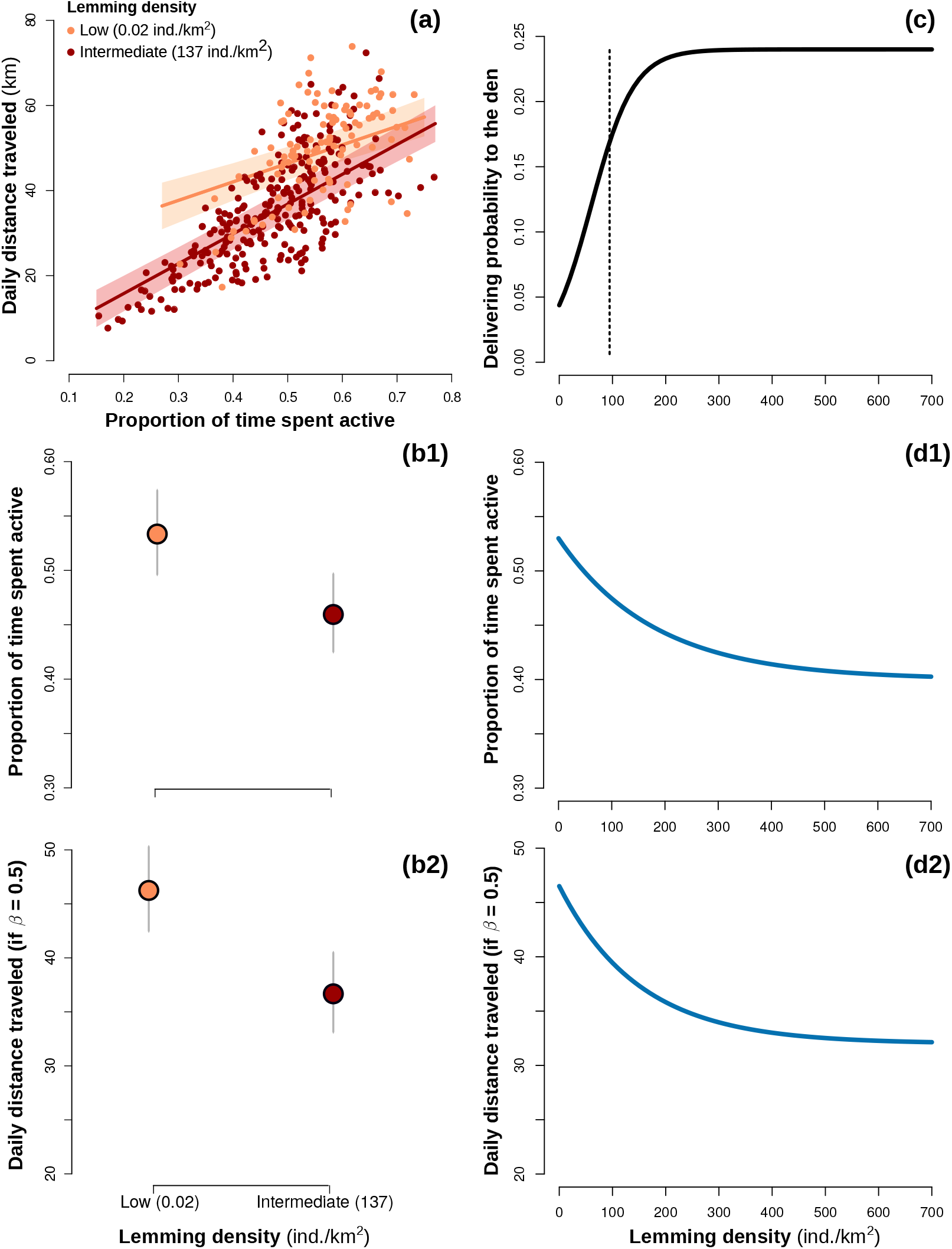
(**a**) Relationship between the daily distance traveled by the arctic fox and the proportion of time it spent active. Colored lines represent predicted relationships and 95% confidence intervals for low and intermediate lemming densities, and dots represent observed data (*n* = 371 fox-days). (**b1**) Predicted proportion of time spent active per day and (**b2**) predicted daily distance traveled by the fox between low and intermediate lemming densities. Error bars represent 95% confidence intervals. Density-dependent relationships of the (**c**) probability that a lemming (prey *i*) is delivered to the den, (**d1**) proportion of time spent active, and (**d2**) daily distance traveled. The dashed line in (**c**) indicates a threshold at which the probability that a fox pair is breeding increases markedly (Juhasz et al., 2020). Figures **c, d1**, and **d2** are partially derived from empirical observations.

The probability that a lemming captured was either consumed, hoarded or delivered to the den was estimated based on behavioral observations of foraging foxes (*n* = 74 in 2004-2005; see Careau et al. 2007). Hoarding and consumption time were estimated with the same method and average 42 sec (*n* = 31) and 32 sec (*n* = 47) respectively. Average delivering time was estimated at 337 sec on the basis that i) dens are generally located close to the centroid of the home range, ii) home ranges average 10 km^2^ (Grenier-Potvin et al., 2021), iii) the average speed of an active fox is 3.8 km h^-1^ (this study), and iv) foxes return an average of 5 lemmings per trip to the den (based on 164 lemming deliveries to the den; Berteaux unpublished data).

Values for the remaining parameters of the functional response of foxes to lemmings, passerines, and sandpipers were extracted from Beardsell et al. (2021) and are summarized in Table 2. Parameter values were estimated using a combination of direct observations of foraging foxes (*n* = 124 hours, 1996-2019), camera traps (2006-2016), and information from the literature (see Beardsell et al. (2021) for more details). We conducted simulations for different values of detection probability for the three prey species since there were a high uncertainty in these parameter values (Beardsell et al. 2021). We presented results of all the simulations in Appendix S2: Figs. S1, S2 and S3. We used a detection and attack probability of 0.15 since the number of lemmings captured per day predicted by the model with this value is more consistent with the highest acquisition rate of foraging foxes observed in the field at high lemming densities (i.e., 2.5 lemming h^-1^ for active foxes; Beardsell et al. 2021). We used intermediate values of 0.12 (*z*_2_) and 0.029 (*z_3_*) in the results.

### 3.4 Density-dependent functions and simulations

We used data from behavioral observations of foraging foxes to define the density-dependent function of the probability that a lemming is delivered to the den. The probability that a lemming captured was delivered to the den is positively related to lemming density (from 0.04 to 0.22 for a year of low and high lemming density respectively; Careau et al. 2007). As the probability that a fox pair is breeding increases markedly around a lemming density of 100 ind. km^-2^ (Juhasz et al., 2020), we used a sigmoidal function to describe the density-dependent relationship between delivering probability and lemming density (Fig. 3c).

We used a combination of foxes accelerometry and GPS tracking data to define the parameter space of the density-dependent functions of the daily proportion of time spent active and distance traveled by the predator. These data were available for two years contrasted by very low (0.02 lemmings km^-2^ in 2018) and intermediate (137 lemmings km^-2^ in 2019) lemming densities. We applied a linear mixed model to estimate the distance traveled and the proportion of time spent active for both lemming densities. We included the distance traveled (km day^-1^) as the response, the proportion of time active and lemming density as fixed effects and individual-fox as random effect. Predicted proportion of time spent active was higher at low than at intermediate lemming density (Fig. 3b1). Predicted daily distance traveled was also higher at low than at intermediate lemming density even when the proportion of active time was set at 0.5 (Fig. 3b2). Based on these results and using the range of values observed for individuals tracked with GPS, as well as the 95% confidence intervals of the average daily distance traveled and the proportion of time spent active recorded over two years, we generated 3 density-dependence functions for each parameter (Figs. 3d1 and d2, Appendix S2: Fig. S1). The model outputs obtained with one function are presented in the results and all other simulations are presented in Appendix S3.

### 3.5 Field evaluation of models

We evaluated the model outputs (A, B, and C) using a 15-year time series (2005-2019) of prey densities and bird nesting success. Lemming densities were estimated annually using live trapping (see Fauteux et al. 2018 for methods). Sandpiper and passerine nest densities were estimated by the maximum number of nests found in an 8 km^2^ plot systematically searched during the nesting season. Each year, nests were revisited every 2-6 days to determine clutch/brood size and nest status (Gauthier et al., 2013; McKinnon et al., 2014). A nest was considered successful if at least one young left the nest (sandpipers) or fledged (passerines). Average annual daily survival rates of passerine and sandpiper nests were estimated using the logistic exposure method (Shaffer, 2004). it was then converted to nest success by increasing daily nest survival to the power of the average number of days (~24 days) between the laying date and the fledging date (for passerines) or hatching date (for sandpipers). See Royer-Boutin (2015) for more details on the calculations of daily nest survival and annual nesting success.

To compare model outputs (A, B and C) to empirical data on bird nesting success, we estimated nesting success of passerines (prey 2) and sandpipers (prey 3) from predator acquisition rates using two differential equations. These equations allow us to calculate predator acquisition rates over the bird nesting period considering that nest density decreases each day. The number of nests predated after 24 days is then divided by the maximum number of nests found in the study plot (*Nb_plot_*), giving us an estimate of annual nesting success. The equation given the total number of passerine nests predated (P_2_) is the product of predator acquisition rate and the number of foxes foraging in the plot (*Nb_fox_*):

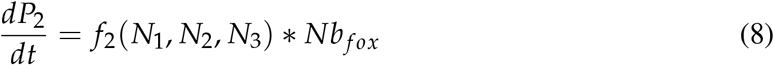

The rate of change in passerine nest density (*N*_2_) is expressed as follows:

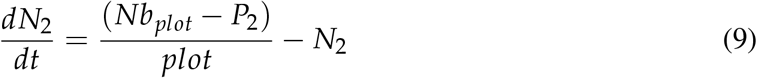

Where *Nb_plot_* is the maximum number of nests found in the study plot, and *plot* the plot size (8 km ^2^). Equivalent equations for sandpiper nests can be obtained by substituting all 2’s for 3’s and vice versa. The model was run for 24 days which corresponds to the duration between the laying date and the fledging date (for passerines) or hatching date (for sandpipers). We assumed that 2 foxes were foraging in the study plot since foxes establish territorial pairs with little overlap between neighboring territories (Clermont et al., 2021; Grenier-Potvin et al., 2021). The model was implemented in R v. 4.0.4 (R Core Team, 2021).

## 4. Results

### 4.1 Multi-prey mechanistic models of functional response

Functional response of the predator (arctic fox) to all prey species were generated for multi-species models with or without density-dependence integrated as explicit functions of prey densities (Fig. 4). According to the Holling traditional multi-species model (model A), the maximum acquisition rate within the range of lemming densities observed in our study system was 30 lemmings·fox^-1^·day^-1^ (Fig. 4a). Including density-dependence in lemmings handling processes through prey delivery (model B) had almost no effect on the functional response of the predator to lemming densities (Fig. 4a; maximum acquisition rate remained the same). Adding density-dependence in predator activity time and distance traveled to the Holling multi-species model (model C) reduced the maximum acquisition rate to 18 lemmings ·fox^-1^·day^-1^ (Fig. 4a). Over the range of prey densities observed in our study system, the functional response of foxes to lemmings does not reach a plateau for all three models. The use of different density-dependence functions of the predator activity time and distance traveled (either linear or hyperbolic) led to relatively similar functional responses of arctic foxes to lemmings (Appendix S3; Fig. S2). Finally, within the range of prey densities observed in our study system, variation in the density of passerine and sandpiper nests had negligible effect on the acquisition rate of lemmings by the predator and, therefore, it is not illustrated in Figure 4a.

**Fig. 4.**
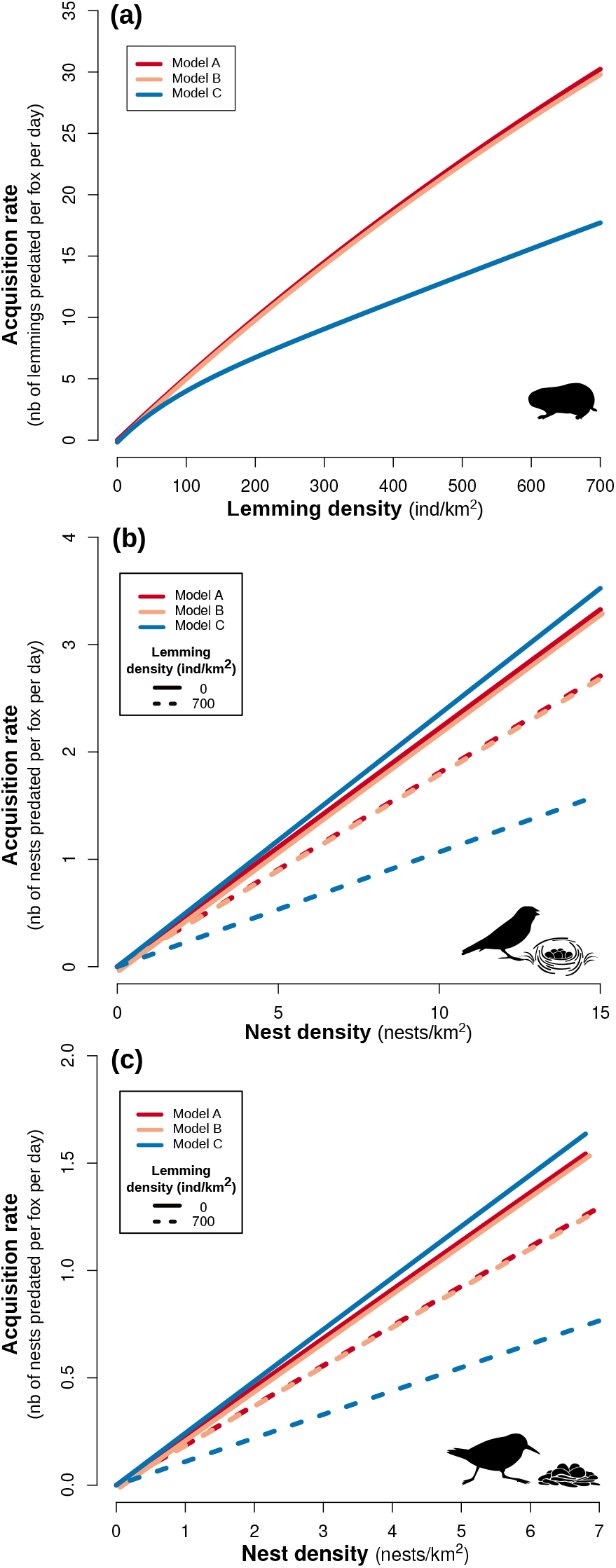
Functional response of the predator (arctic fox) to prey 1 (lemmings; **a**), prey 2 (passerine nests; **b**) and prey 3 densities (sandpiper nests; **c**) following models A, B, and C (each colored line represents a model). Model A is the Holling multi-species model, model B modifies the Holling multi-species model by adding density-dependence in lemmings handling processes through prey delivery, and C modifies the Holling multi-species model by adding density-dependence in predator activity time and distance traveled. Densities of prey 2 and 3 are set at intermediate densities in **a**.

Acquisition rate of passerine and sandpiper nests by the predator decreased slightly with increasing lemming density when considering only prey handling processes in the multi-species model without density-dependence (model A). The slope of acquisition rate was 18% lower when comparing a low (0 ind. km^-2^) and a high (700 ind. km^-2^) lemming year (Figs. 4b and 4c). Similarly, including density-dependence in lemming handling processes (model B) reduced the slope of nests acquisition rate by 19% between a low and high lemming year. Finally, as illustrated in Figs. 4b and 4c, the models including density-dependence in predator activity time and distance traveled to the Holling multi-species model (model C) had the most pronounced effect as it reduced the slope of passerine and sandpiper nest acquisition rate by 54% between a low and high lemming year.

### 4.2 Field evaluation of the multi-prey models

Summer lemming density varied from 2 to 648 individuals per km^2^ from 2005 to 2019. A total of 625 passerine nests were monitored during this period and annual nesting success of passerines averaged 48% (range: 8 to 88%; Fig. 5a). From 2005 to 2019, 292 sandpiper nests were monitored and annual nesting success of sandpipers averaged 50% (range: 4 to 100%; Fig. 5b).

**Fig. 5.**
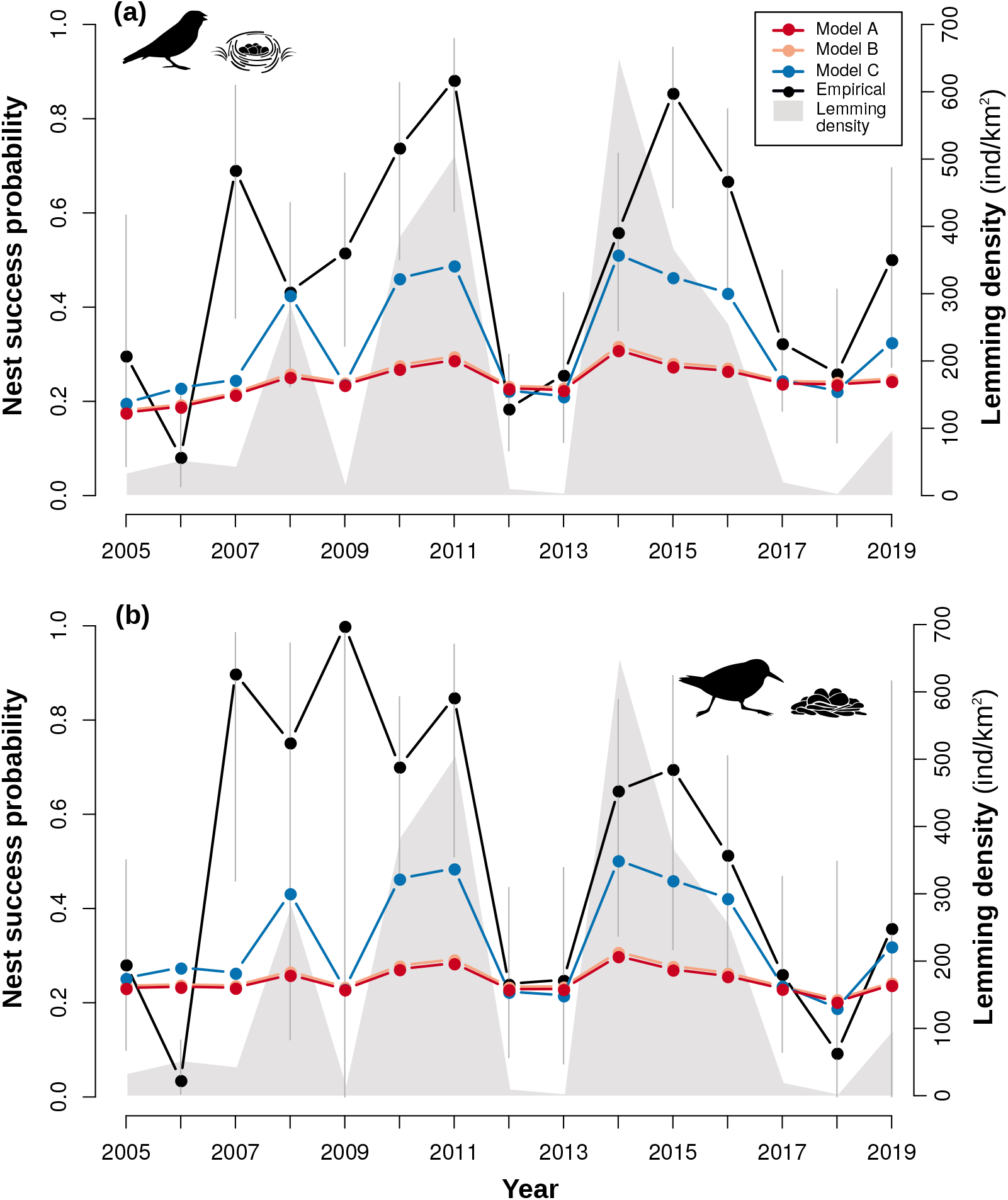
Temporal fluctuations in lemming densities (cyclic prey 1), and nesting success of passerines (**a**; prey 2) and sandpipers (**b**; prey 3). Model results and empirical time series are represented by a different color. Model A is the Holling multi-species model, model B modifies the Holling multi-species model by adding density-dependence in lemmings handling processes through prey delivery, and C modifies the Holling multi-species model by adding density-dependence in predator activity time and distance traveled. Model results for A and B overlap. Empirical data (lemming density and birds nest success) were acquired through long-term monitoring in the Canadian High-Arctic. Error bars are the 95% confidence intervals of nesting success estimates.

When considering only prey handling processes, model A and B generated very limited temporal variations in bird nesting success (Fig. 5). This indicates that lemming handling time alone cannot generate fluctuations in annual bird nesting success consistent with our time series. Model C including density-dependence in predator foraging behavior generated temporal variations in bird nesting success that were relatively consistent with those observed in our study system, including a marked release in predation pressure at high lemming densities (Fig. 5). However, annual variations in bird nesting success generated by model C were of smaller amplitudes than empirical observations. Our main results were robust to variation in density-dependent functions of the predator daily proportion of time spent active and distance traveled (see Appendix S3).

## 5. Discussion

In this study, we first derived a mechanistic multi-species functional response model by breaking down key components of predation (i.e., search, attack, pursuit, handling). We also incorporated prey density-dependence in predator foraging behavior into the Holling multi-species model. We then applied this model to an intensively studied arctic vertebrate community to evaluate the relevance of various proximate mechanisms that could explain the short-term positive indirect effects observed between tundra prey species. We showed that handling processes cannot explain the predation release on nesting birds observed during lemming peaks. However, adding prey density-dependence in arctic fox foraging behavior generated variation in nest predation rate consistent with long–term empirical data. Such a mechanism has been little studied to date and may play a significant role in modulating species interaction rates. To our knowledge, our study is the first attempt to disentangle key components of predation in a natural multi-prey system, and we think that this approach could be applied to a broad range of food webs.

Traditional, multi-species quantitative models are commonly used and assume a saturation of predator acquisition rates with increasing prey availability due to rate-limiting handling processes (Turchin and Hanski, 1997; Matthiopoulos et al., 2007; McLellan et al., 2010; Serrouya et al., 2015). Beyond the consequences of assuming predator saturation in population dynamics models, this assumption may have consequences on proposed management approaches (Griffen, 2021). For instance, one approach recently proposed was to feed a mammalian predator to reach saturation more quickly, and hence reduce predation pressure on a threatened prey species (Spencer et al., 2016). However, predator saturation by prey handling processes does not appear to be frequently observed in the wild (Novak, 2010; Chan et al., 2017; Preston et al., 2018), and our results showed that fundamentally different mechanisms may limit predator acquisition rate. Handling time is often estimated by fitting a statistical functional response model to empirical data (Smout et al., 2010; Paterson et al., 2015) but as pointed out by Griffen (2021), this method does not ensure that handling time is an ecologically meaningful parameter. Thus, we emphasize the need for a mechanistic understanding of functional response and, when traditional statistical models (Holling’s type II and III) are fitted, care must be taken in interpreting predator foraging behavior.

In our study, we extended the Holling multi-species model by adding density-dependence in prey handling processes through prey delivery to offspring. While this mechanism plays a minor role in our study system, the sensitivity of predator acquisition rate to this modification of the activity budget is likely to depend on predator home range size, prey load-size, predator movement rate and predator ability to forage while delivering food. For instance, prey delivery could represent a significant proportion of the activity budget for a predator with a large foraging range (e.g., albatrosses; Weimerskirch et al. 1993) or constrained to bring one prey at a time to the breeding site. This type of prey density-dependent mechanism remains to be explored in other natural predator-prey systems.

Numerous studies highlight the relevance of improving functional response models and integrating alternative mechanisms that can modulate animal acquisition rates, such as predator activity level (Toscano and Griffen, 2014), prey digestion time (Jeschke, 2007; Papanikolaou et al., 2020) or time spent in vigilance (Sirot et al., 2021). We found evidence that changes in predator foraging behavior with respect to main prey density can at least partly explain the positive indirect effects observed in a vertebrate community. Although many empirical studies have demonstrated links between prey availability and predator foraging behavior (e.g., Harding et al. 2007; Bertrand et al. 2014; Busdieker et al. 2019), dependence of predator foraging behavior on prey density is rarely included in predator-prey models (but see Abrams 1982). We recognize that the density-dependent functions used in our study were derived from limited empirical data. Further field investigations, such as long-term GPS and accelerometer tracking of predators over a range of prey densities, are needed to refine functions and fully integrate density-dependent changes in predator foraging behavior. Overall, our results highlight the need to reinforce the links between multi-species functional response models and the dynamics of vertebrate communities.

The short-term, positive indirect effects of lemmings on tundra nesting birds due to shared predators were reported several decades ago and studied over the circumpolar arctic (Underhill et al., 1993; Summers et al., 1998). Various mechanisms have been proposed, including predator satiation and prey switching, but these mechanisms have never been demonstrated (Underhill et al., 1993; Summers et al., 1998; Blomqvist et al., 2002; McKinnon et al., 2014; Bowler et al., 2020). Our results indicate that predation release on tundra birds at high lemming densities is primarily due to a reduction in arctic fox daily activity time and distance traveled. Prey switching is another mechanism that has been suggested to explain predation release (Blomqvist et al., 2002; Bêty et al., 2002; Iles et al., 2013; Bowler et al., 2020). Prey switching refers to a situation where the preference for a prey *i* by the predator is greater when prey *i* is abundant relative to other prey, and inversely smaller when prey *i* is less abundant than other prey (Murdoch, 1969). An increase in preference for an abundant prey *i* may translate into an increase in the probability of its detection, attack and/or success as prey *i* density increases because of changes in predator behavior or foraging strategy.

Prey switching remains to be fully explored in our study system and more empirical data are needed to investigate the effect of lemming density on the probability of lemming and bird nest detection by foxes, as well as their attack and success probability. However, this mechanism is expected to play a relatively minor role. Indeed, even if foxes capture more lemmings when they are abundant, the handling time per lemming captured is still likely to be too low to have a significant effect on bird nest predation rates. Moreover, highly vulnerable prey like passerines and sandpipers are unable to protect their clutches against arctic foxes (Smith and Edwards, 2018; Hussell and Montgomérie, 2020). Consequently, once the nest is detected, the probability of nest attack by foxes is likely to remain high in all years as attacking these nests engenders very low costs (i.e., low handling time and no risk of injury) but systematically provides benefits to the predators. Annual changes in the probability of nest attack by foxes is however more likely in large-bodied tundra nesting birds, as reported in snow geese (Bêty et al., 2002). Our multi-species model could be adapted to explore the effects of such changes on the nesting success of bird species able to protect their clutches, fight back and defeat arctic foxes.

Some variability in shorebird and passerine nesting success remains unexplained in our study system and this can be the result of a combination of factors. First, we assumed that lemming density and parameter values were homogeneous across the landscape. Better knowledge of such variation, especially within fox territories, would likely contribute to explaining variation in bird nesting success. Second, the empirical measurement of nesting success may be overestimated since nests predated very early in the nesting period were not necessarily found by observers. Third, in some years, nesting success was estimated from a limited number of nests (as indicated by the wide confidence intervals in Fig. 5). Finally, we assumed that two foxes were foraging in the monitoring area of passerine and sandpiper nests in all years. Although this is most likely the predominant situation (Clermont et al., 2021), fox number is likely to generate variation in bird nesting success since territories may overlap and/or be occupied by three foxes in some years (6 cases, *n*=130 social units; Lai 2017).

The effect of a predator on prey population dynamics will not only depend on the functional response, but also on the numerical response of the predator to changes in density of all prey species. Our analysis here focused on per capita impact of predators. A full exploration of the implications of the per population impact is beyond the scope of this paper and requires further analysis of fox population dynamics. Within the fox-passerine-sandpiper system, numerical response does not appear to be an important driver as the number of territorial adult foxes remains relatively constant from one summer to another, even if the breeding success of foxes is strongly influenced by lemming cycles (Royer-Boutin, 2015; Juhasz et al., 2020). However, the presence of another abundant prey species in the system, like the colonial nesting snow goose, can increase predation pressure on passerines and sandpipers (McKinnon et al., 2014; Flemming et al., 2016) likely through an aggregative numerical response by foxes (Lamarre et al., 2017). The combined effect of lemming cycles and goose presence on the functional and numerical response of foxes, and its impact on passerine and sandpiper nesting success, deserve further investigations.

In this study, we extended the Holling multi-species model to incorporate key ecological processes. We hope that our work will provide a useful starting point to build mechanistic models of predation, adapted to natural systems like vertebrate communities. Although the model was inspired by an active hunting predator, it should stimulate further studies, including mechanistic models that integrate ecological processes and constraints relevant to various multi-species system (e.g., predator hunting strategy, predator interference and group formation). In recent years, there has been a growing number of studies that aim to predict trophic links based on species traits, especially body size (Gravel et al., 2013; Ho et al., 2019; Portalier et al., 2019) but mechanistic models quantifying interaction strength in a multi-species context are still lacking. With recent advances in biologging technology, high-frequency GPS, acoustic and accelerometer data are increasingly used to study free-ranging organisms (Williams et al., 2014; Pagano et al., 2018; Studd et al., 2019). The parametrization of mechanistic models with such data is a promising method to quantify interaction strength in natural systems (Merrill et al., 2010; Noonan et al., 2021; Studd et al., 2021).

## 6. Acknowledgments

We thank Mark Novak for helpful comments and suggestions on this manuscript. We also thank Éliane Duchesne for her contribution to the analyses of bird nesting success data and Madeleine-Zoé Corbeil-Robitaille for the illustrations. The research relied on the logistic assistance of the Polar Continental Shelf Program (Natural Resources Canada) and of Sirmilik National Park of Canada. The research was funded by (alphabetical order): Arctic Goose Joint Venture, the Canada Foundation for Innovation, the Canada Research Chairs Program, the Canadian Wildlife Service (Environment Canada), the Fonds de recherche du Québec-Nature et technologies, the International Polar Year program of Indian and Northern Affairs Canada, the Natural Sciences and Engineering Research Council of Canada, the ArcticNet Network of Centers of Excellence, the Northern Scientific Training Program, Polar Knowledge Canada, Université du Québec à Rimouski, Université Laval and the W. Garfield Weston Foundation. Finally, we are especially grateful to the many people who helped us with field work over many years, the Mittimatalik Hunters and Trappers Organization and Park Canada’s staff for their assistance.

## 7. Appendix S1: Derivation of the general multi-species functional response model

The area searched (*A;* km^2^) by the predator is expressed by the product of the distance traveled per day when the predator is active (*s*, km/day), the reaction distance to a prey item *i* (*d_i_*, km), and the time spent searching (*T_s_*, day):

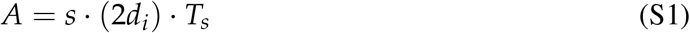

A potential encounter occurs between the predator and a prey item *i* when the predator is at a distance (d_i_), being defined as the maximum distance at which predator and prey can detect or react to each other (in 2D, detection region = 2*d*; Pawar et al. 2012). As not all prey within this area may be detected, attacked and subdued by the predator, we introduced the detection probability (*z_i_*), the attack probability (*k_i_*), and the success probability of an attack (*p_i_*). Capture efficiency of a prey item *i* (*α_i_*, km^2^/day) by the predator is expressed by:

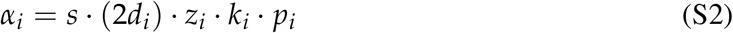

The number of prey captured (*V_αi_*) during the time spent searching *T_S_* (day) for a density *N_i_* (number of *i*/km^2^) is:

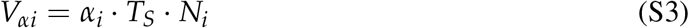

The time spent searching (*T_S_*) is partitioned into three mutually exclusive components:

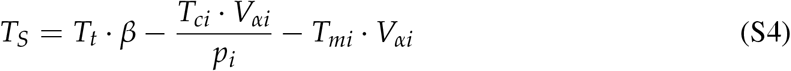

where *T_t_* is the total time available in a day which is reduced by i) *β* which expresses the proportion of time spent active in a day by ii) 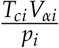 which is the time spent chasing a prey *i* once encountered and by iii) *T_mi_V_αi_* which is the time spent handling a prey *i* if subdued. Note that we can combine the chasing and manipulation time to produce an overall prey handling time (*h_i_*, day/*i*) of the prey *i*:

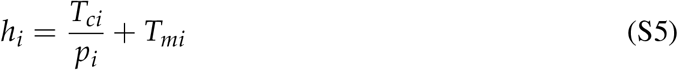

Equivalent equations (S1 to S5) of a prey of species *j* and *k* can be obtained by substituting all *i*’s for *j*’s (or *k*). In order to extend the model to multiple prey species (here *i*, *j* and *k*), we also substract the time spent chasing and manipulating a prey *j* or *k*:

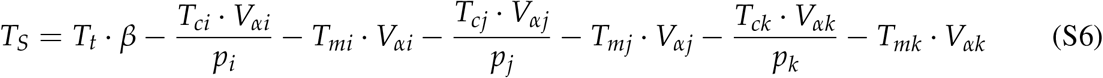

When searching, the predator is not looking for one specific prey; the time spent searching is common to all prey. By simplifying Eq. S6 we have:

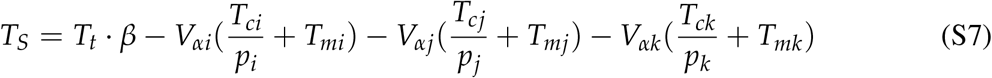

Substituting *T_s_* from Eq. S7 into Eq. S3, we arrive at:

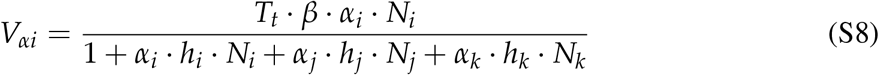

The functional response of a predator to a prey species *i* is the number of prey captured per predator per unit of time (*f_i_* (*N_i_*, *N_j_*, *N_k_*)). This is expressed by dividing Eq. S8 by T_t_:

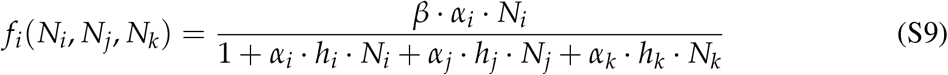

An equivalent equation for the predator’s acquisition rate on prey *j* and *k* can be obtained by substituting all *i*’s for *j*’s (or *k*) in equation S9 and vice versa.

## 8. Appendix S2: Simulations of detection and attack probability values

**Fig. S1.**
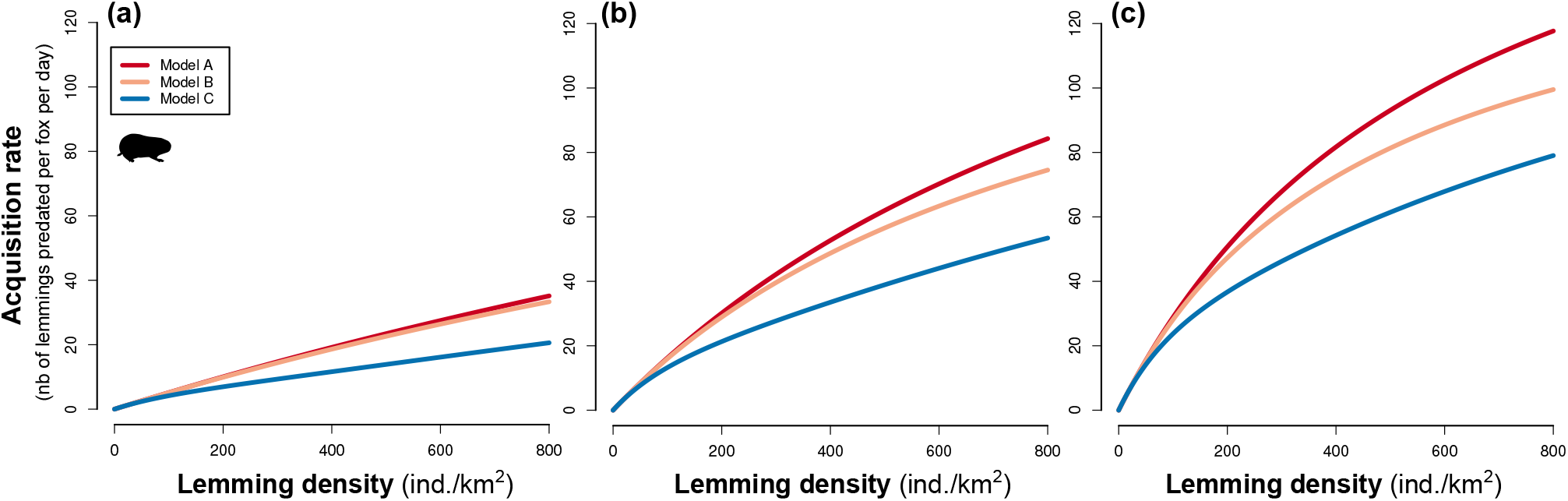
Functional response of the predator (arctic fox) to prey 1 density (lemming) for 3 values of the detection and attack probability of a lemming by the fox within the maximum reaction distance (**(a)**: *z*_1_ * *k*_1_ = 0.15, **(b)**: *z*_1_ * *k*_1_ = 0.50, and **(c)**: *z*_1_ * *k*_1_ = 0.95). Each model is represented with a different color.

**Fig. S2.**
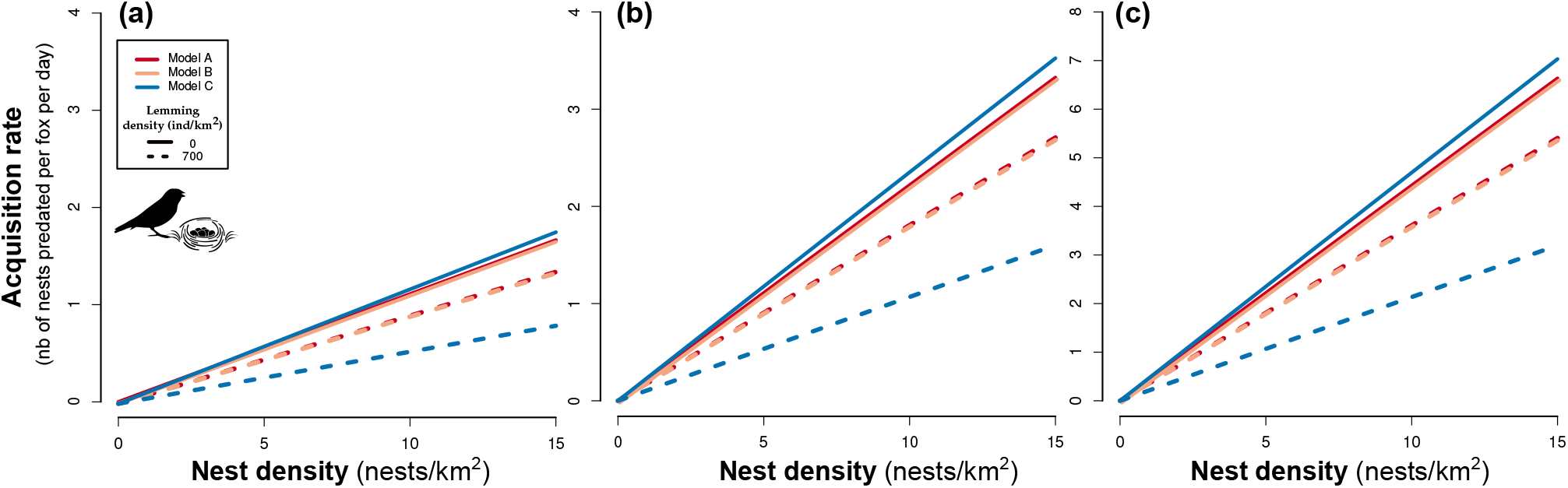
Functional response of the predator (arctic fox) to prey 2 density (passerine nests) for 3 values of the detection probability of a passerine nest by the fox within the maximum reaction distance ((a): *z*_2_ = 0.06, (b): *z*_2_ = 0.12, and (c): *z*_2_ = 0.24). Each model is represented by a different color at two lemming densities.

**Fig. S3.**
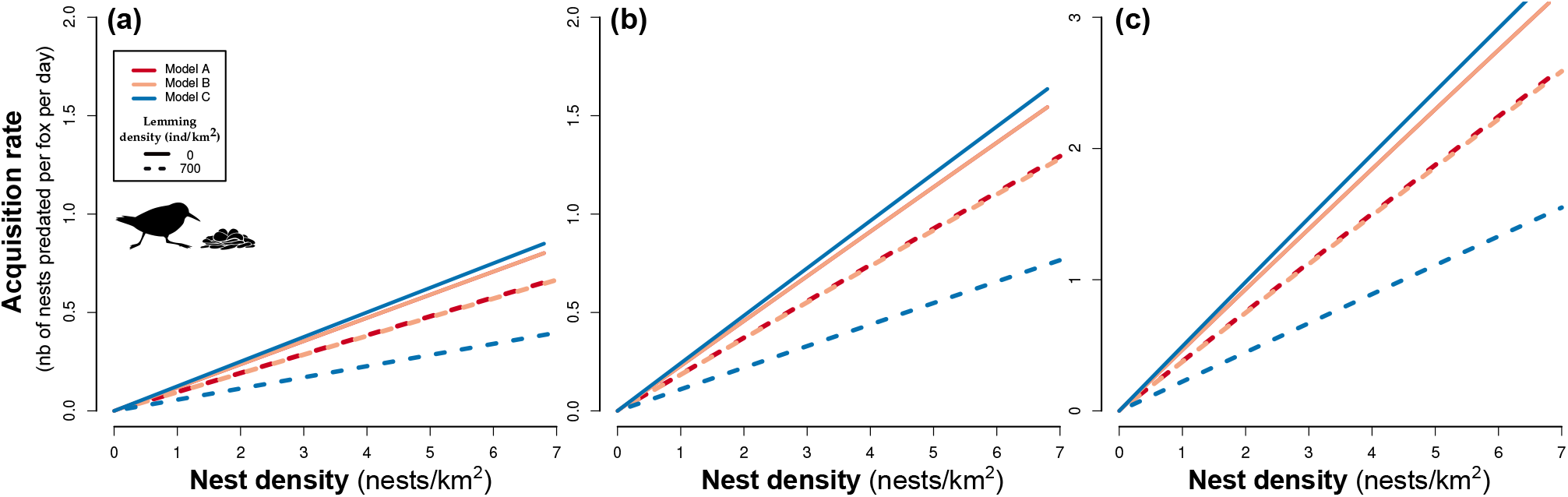
Functional response of the predator (arctic fox) to prey 3 density (sandpiper nests) for 3 values of the detection probability of a sandpiper nest by the fox within the maximum reaction distance ((a): *z*_3_ = 0.015, (b): *z*_3_ = 0.029, and (c): *z*_3_ = 0.059). Each model is represented by a different color at two lemming densities.

## 9. Appendix S3: Simulations of density-dependence functions of the predator activity time and distance traveled

**Fig. S1.**
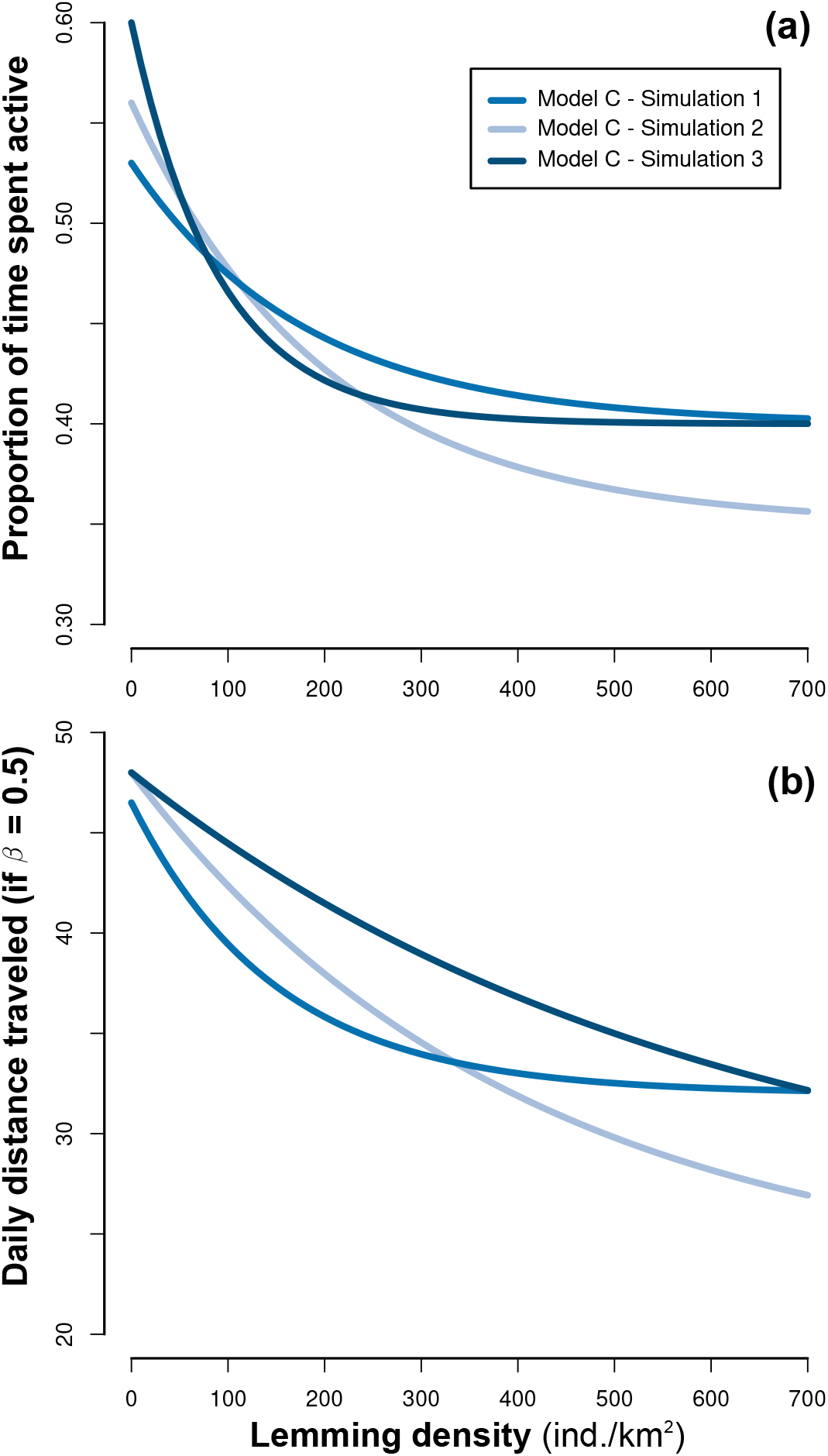
Simulations of three density-dependence functions of the predator daily proportion of time spent active (**a**) and distance traveled (**b**). Each colored line represents a simulation.

**Fig. S2.**
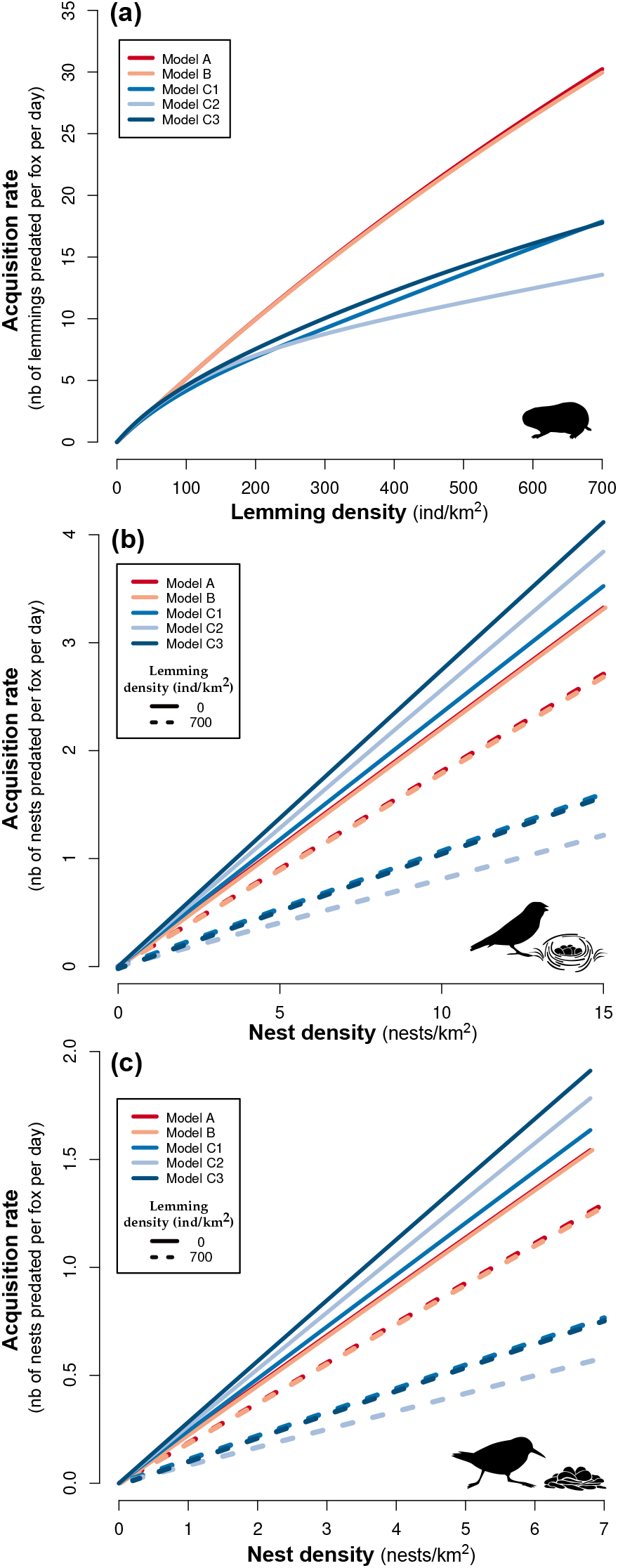
Functional response of the predator (arctic fox) to prey 1 (lemmings; **a**), prey 2 (passerine nests; **b**) and prey 3 densities (sandpiper nests; **c**) following models A, B, and C (each colored line represents a model or a simulation). Model A is the Holling multi-species model, model B modifies the Holling multi-species model by adding density-dependence in lemmings handling processes through prey delivery, and C modifies the Holling multi-species model by adding density-dependence in predator activity time and distance traveled. Densities of prey 2 and 3 are set at intermediate densities in **a**.

**Fig. S3.**
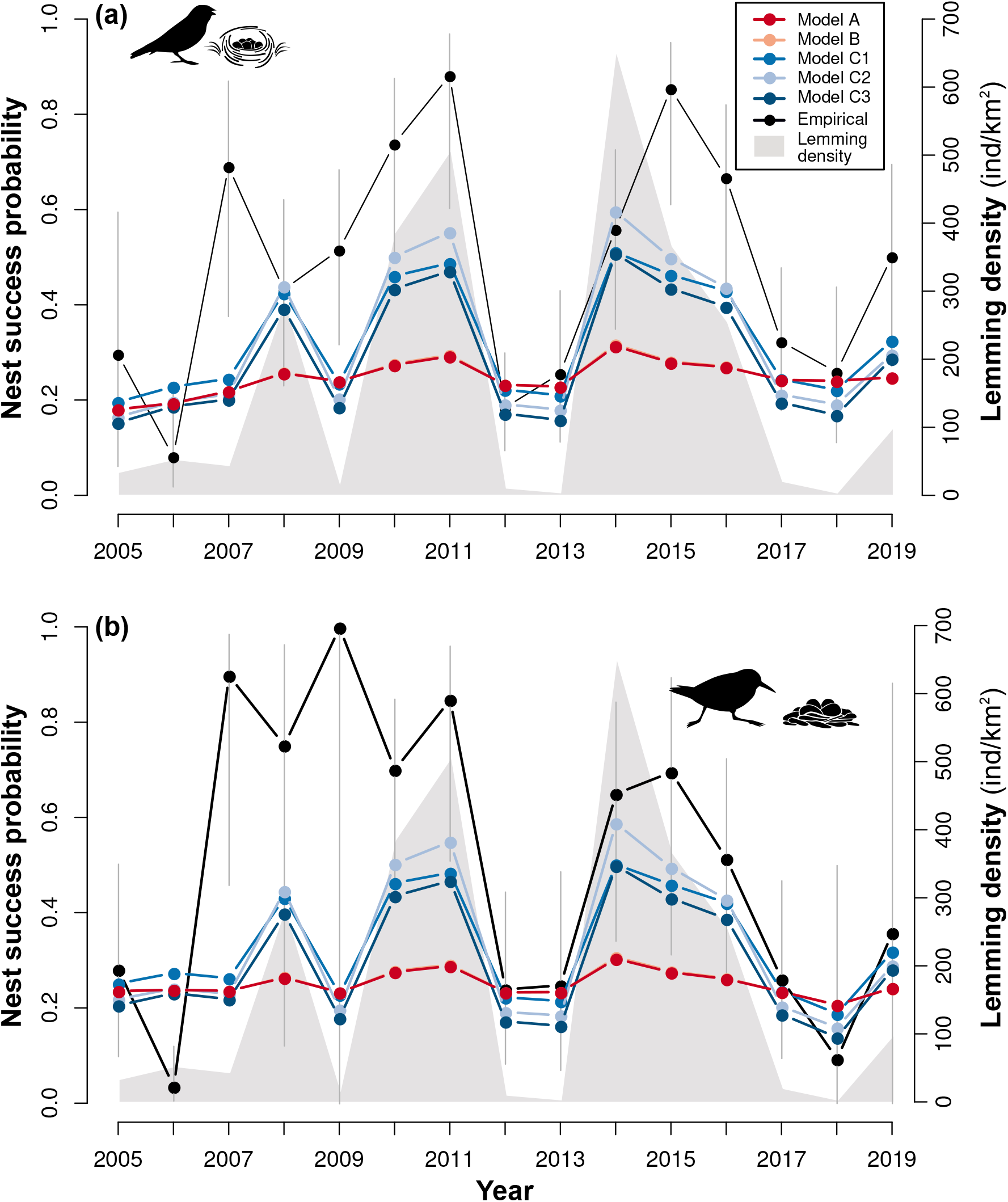
Temporal fluctuations in lemming densities (cyclic prey 1), and nesting success of passerines (**a**; prey 2) and sandpipers (**b**; prey 3). Model results and empirical time series are represented by a different color. Model A is the Holling multi-species model, model B modifies the Holling multi-species model by adding density-dependence in lemmings handling processes through prey delivery, and C modifies the Holling multi-species model by adding density-dependence in predator activity time and distance traveled. Model results for A and B overlap. Empirical data (lemming density and birds nest success) were acquired through long–term monitoring in the Canadian High-Arctic. Error bars are the 95% confidence intervals of nesting success estimates.

